# Developmental remodeling repurposes larval neurons for sexual behaviors in adult *Drosophila*

**DOI:** 10.1101/2023.11.30.569472

**Authors:** Julia A. Diamandi, Julia C. Duckhorn, Kara E. Miller, Mason Weinstock, Sofia Leone, Micaela R. Murphy, Troy R. Shirangi

**Affiliations:** Villanova University, Department of Biology, 800 East Lancaster Ave, Villanova, PA 19085 U.S.A.

## Abstract

Most larval neurons in *Drosophila* are repurposed during metamorphosis for functions in adult life, but their contribution to the neural circuits for sexually dimorphic behaviors is unknown. Here, we identify two interneurons in the nerve cord of adult *Drosophila* females that control ovipositor extrusion, a courtship rejection behavior performed by mated females. We show that these two neurons are present in the nerve cord of larvae as mature, sexually monomorphic interneurons. During pupal development, they acquire the expression of the sexual differentiation gene, *doublesex*, undergo *doublesex*-dependent programmed cell death in males, and are remodeled in females for functions in female mating behavior. Our results demonstrate that the neural circuits for courtship in *Drosophila* are built in part using neurons that are sexually reprogrammed from former sex-shared activities in larval life.

## INTRODUCTION

Many behaviors of adult animals develop during a period of maturation, *e.g.*, puberty, as juveniles transform into adults^1^. The central nervous system changes dramatically during this time, including the birth of new neurons that build adult circuits^2^. However, in some animals like insects^3^ and worms^4^, the adult nervous system is also assembled with reprogrammed neurons that were formerly active in the juvenile.

During *Drosophila* metamorphosis, most neurons of the larval central nervous system are recycled for use in adult circuits^5^. The moonwalker descending neuron, for instance, triggers backward locomotion in crawling larvae and is remodeled during pupal life to regulate backward walking in adult flies^6^. Additionally, several input and output neurons of the larval mushroom body trans-differentiate during metamorphosis and contribute to entirely different circuits in the adult brain^3^. Despite these cases, there are currently no examples of recycled larval neurons that contribute to sexually dimorphic behaviors of adult flies. Indeed, lineage analyses of adult neurons with sexual identity in *Drosophila* suggest that most are born post-embryonically and contribute exclusively to sexually dimorphic behaviors in adults^7,8^.

We previously identified a small sexually dimorphic population of interneurons in the abdominal ganglion of adult flies, the DDAG neurons, that co-express the *tailless*-like orphan nuclear receptor, *dissatisfaction* (*dsf*), and the sex differentiation gene, *doublesex* (*dsx*)^9^. Here, we identify two female-specific DDAG neurons that influence the extrusion of the ovipositor performed by mated females to reject courting males. These two DDAG neurons are born during embryogenesis and exist as segmental homologs of a sexually monomorphic interneuron in the larval abdominal ganglion called A26g. During early pupal life, the A26g neuron at abdominal segments five and six acquire the expression of *dsx*, undergo *dsx*-dependent programmed cell death in males, and are remodeled in females for use in the circuitry for ovipositor extrusion. Our results demonstrate that the neural circuits for sexually dimorphic behaviors in *Drosophila* include sexually reprogrammed neurons with former activities in the juvenile larva of both sexes.

## RESULTS

### The DDAG neurons are anatomically diverse

*Drosophila* (*D.*) *melanogaster* females and males have eleven and three *dsf*- and *dsx*-co-expressing abdominal ganglion (DDAG) neurons, respectively, which contribute to several female- and male-specific mating behaviors (Figure 1A)^9^. As a first step toward associating specific courtship functions to specific DDAG neurons, we employed a stochastic labeling method to determine the anatomy of individual DDAG neurons in females and males.

**Figure 1.**
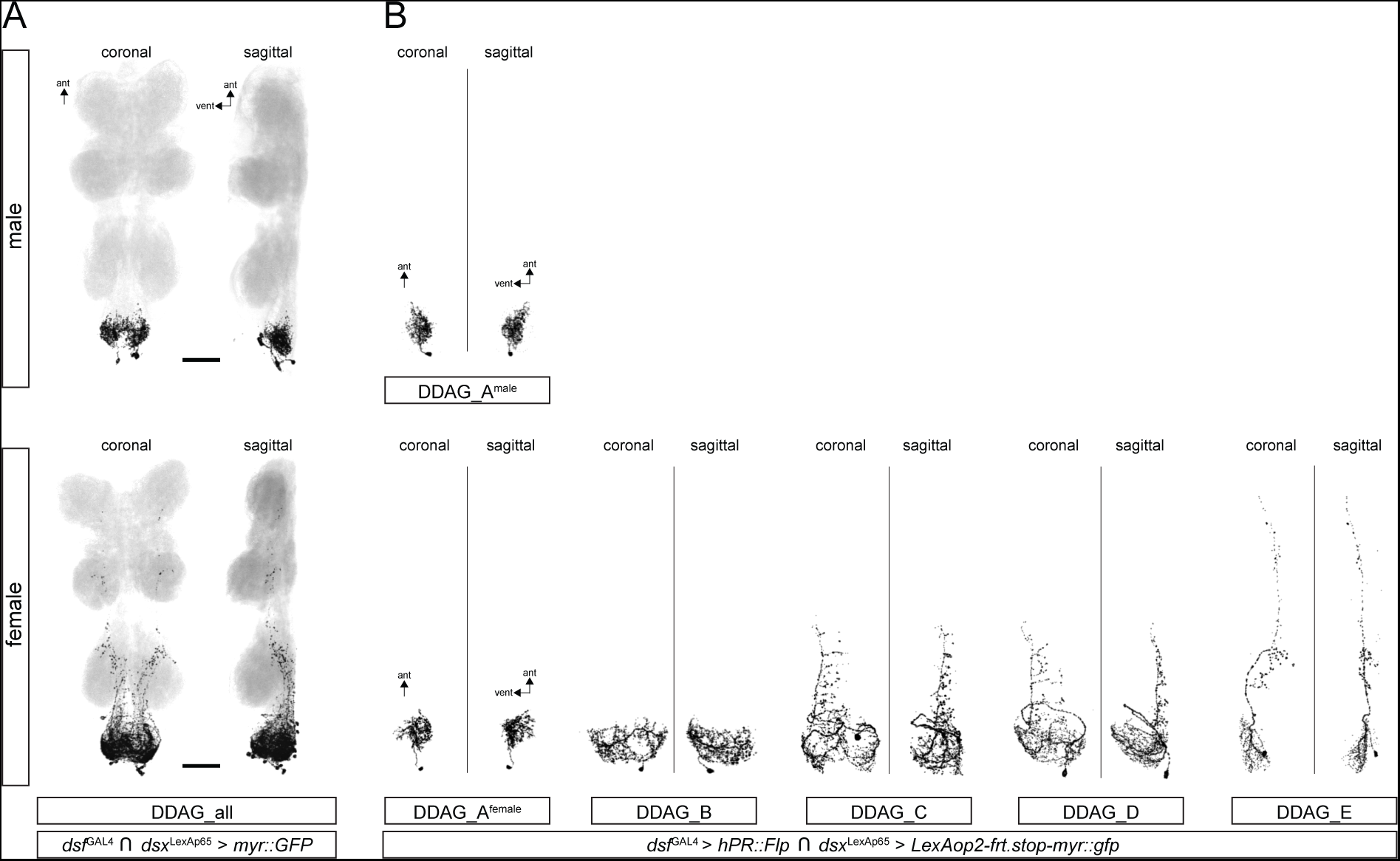
The DDAG neurons are anatomically diverse. (A) Confocal images of ventral nerve cords (VNCs) from *dsf*^Gal4^ ∩ *dsx*^LexA::p65^ > *myr::gfp* males and females. GFP-expressing neurons are labeled in black and DNCad (neuropil) is shown in light gray. (B) Confocal images of individual DDAG neurons from *dsf*^Gal4^/*LexAop-frt.stop-myr::gfp*; *dsx*^LexA::p65^/*UAS-hPR::Flp* males and females fed food containing mifepristone for about two hours during adulthood. A total of one and five DDAG subtypes were identified in males and females, respectively. Coronal and sagiaal views of each neuron is shown.

The DDAG neurons are labeled by a genetic intersectional strategy whereby a Flp recombinase, driven in *dsx*-expressing cells by *dsx*^LexA::p65^ ^10^, excises a transcriptional stop casseae from an upstream activating sequence (UAS)-regulated *myr::gfp* transgene. Expression of *myr::gfp* is activated in *dsf*-co-expressing cells by the *dsf*^Gal4^ allele^9^. To stochastically label individual DDAG neurons using a similar intersectional strategy, we utilized *FlpSwitch*^11^, a Flp recombinase fused to the ligand binding domain of the human Progesterone Receptor (hPR). The activity of the FlpSwitch recombinase is dependent upon the presence of the progesterone mimic, mifepristone. We constructed flies carrying the *dsf*^Gal4^ and *dsx*^LexA::p65^ alleles, a UAS-regulated *FlpSwitch*, and a LexAop-regulated *myr::gfp* transgene containing a transcriptional stop casseae that is conditionally excised by the FlpSwitch recombinase. When these flies were fed food containing mifepristone for a relatively short period of time, single DDAG neurons were often labeled in adult flies. We examined approximately 60 and 20 single cell clones in female and male ventral nerve cords, respectively, and identified five anatomically distinct DDAG subtypes (Figure 1B). One of these subtypes, DDAG_A, is present in both sexes, whereas the DDAG_B–E subtypes are specific to females. All DDAG neuronal subtypes arborize extensively in the abdominal ganglion, whereas the DDAG_C–E neurons also extend an anteriorly projecting branch that innervates thoracic neuropils. We conclude that the DDAG neuronal population consists minimally of five anatomical subtypes. Additional DDAG subtypes may exist that were not labeled in our experiments, and the number of neurons of each subtype is unclear.

### DDAG_C and DDAG_D neurons contribute to ovipositor extrusion

The five DDAG subtypes we identified in females may contribute to different reproductive behaviors. To develop driver lines that target subsets of DDAG neurons, CRISPR/Cas9-mediated homology-directed repair was used to create new alleles of *dsf* that express *LexA::p65*, or the Split-Gal4 hemi-drivers, *p65AD::Zp* and *Zp::GAL4DBD*, in *dsf*-expressing cells. In a search for Split-Gal4 drivers that target subsets of DDAG neurons, we found that intersecting *dsf*^p65AD::Zp^ with the enhancer line, *VT026005*-*Zp::GDBD*, labeled four female-specific DDAG neurons per hemisphere of the adult ventral nerve cord (Figure 2A, B). Two cell bodies are located at the posterior tip of the nerve cord and the other two are at the dorsal side of the abdominal ganglion. Labeling of individual neurons by the multicolor Flp-out technique^12^ identified the two neurons at the tip as a local interneuron corresponding to the DDAG_B subtype, whereas the two dorsally located cell bodies are the DDAG_C and DDAG_D neurons (Figure 2C). The DDAG_C and DDAG_D neurons are segmental homologs (see below) with several anatomical similarities (Video S1). Both neurons extend a primary branch off the cell body forming a “ventral arch” that projects across the midline and then anteriorly on the contralateral dorsal side. Unlike the DDAG_D neuron, the DDAG_C neuron extends a dorsally projecting medial branch off the ventral arch that gives rise to arbors within the abdominal ganglion. The DDAG_D neuron is located anterior to the DDAG_C neuron. We were unable to differentiate the two DDAG_B neurons, however they may exhibit subtle anatomical differences that were undetected in our analysis and may contribute to female behavior differently.

**Figure 2.**
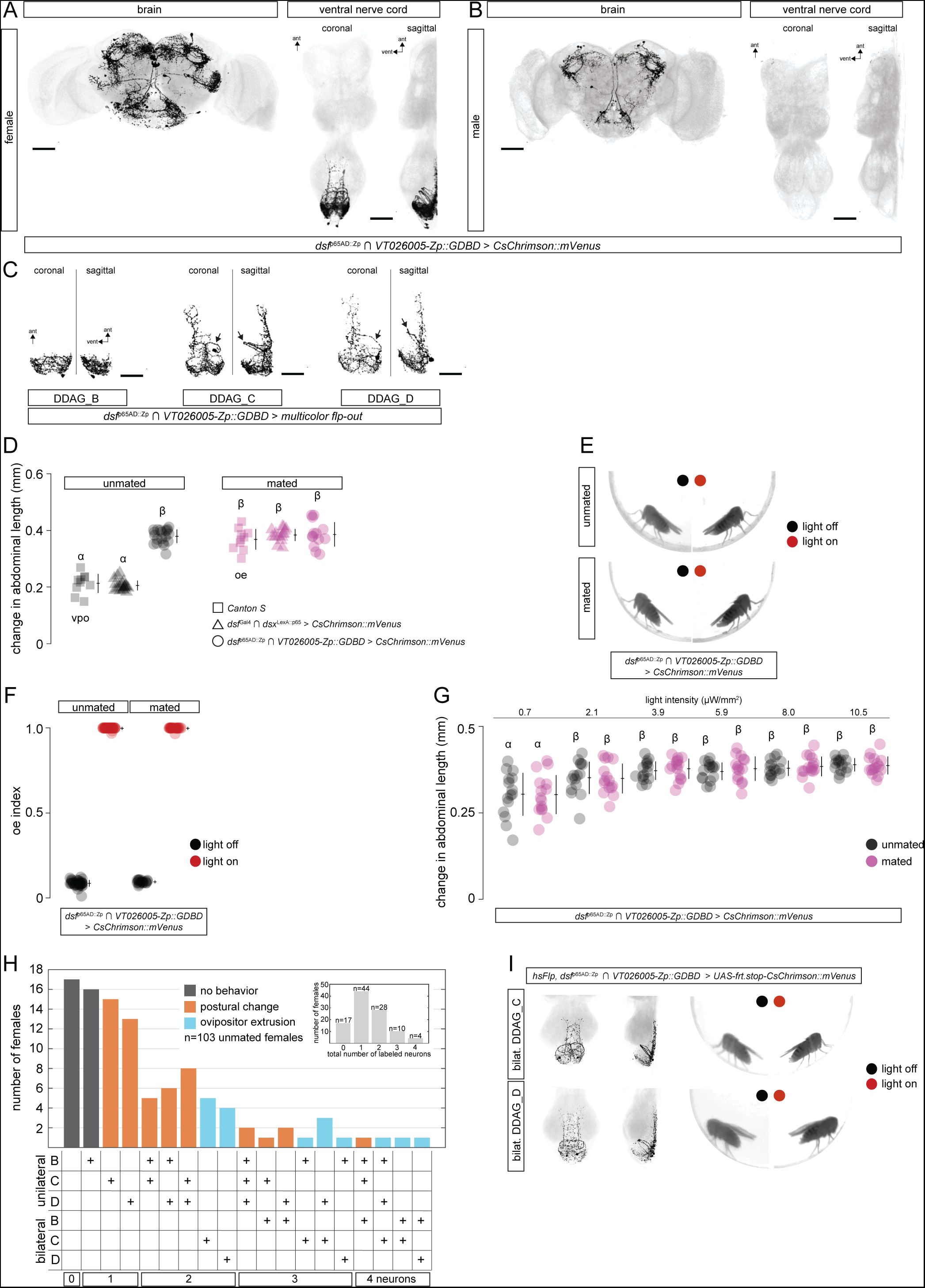
The DDAG_C and DDAG_D neurons contribute to ovipositor extrusion in unmated and mated females. (A and B) Confocal images of brains and VNCs from *dsf*^p65AD::Zp^ ∩ *VT026005-ZP::GDBD* > *CsChrimson::mVenus* females and males probed with antibodies to GFP. *CsChrimson::mVenus*-expressing neurons are labeled in black and DNCad (neuropil) is shown in light gray. Four VNC neurons are labeled per hemisphere. (C) Confocal images of individual DDAG_B, DDAG_C, and DDAG_D neurons from *dsf*^p65AD::Zp^ ∩ *VT026005-ZP::GDBD* > multicolor flp-out females. Arrows point to the ventral arch of the DDAG_C and DDAG_D neurons. The DDAG_C neuron has a medial branch off the ventral arch that is absent in the DDAG_D neuron. (D) Artificial activation of the DDAG_B–D neurons induces ovipositor extrusion in unmated and mated females. The change in abdominal length when an unmated (black squares) or mated (magenta squares) *Canton S* female performs vaginal plate opening or ovipositor extrusion, respectively, during courtship is shown. Photoactivation of the entire DDAG population in unmated (black triangles) and mated (magenta triangles) females induces a change in abdominal length like that observed when *Canton S* females open their vaginal plates or extrude their ovipositor, respectively^9^. Photoactivation of the DDAG_B–D subtypes in unmated (black circles) and mated (magenta circles) females induces a change in abdominal length like that observed when *Canton S* mated females extrude their ovipositor. (E) Still-frame images of decapitated *dsf*^p65AD::Zp^ ∩ *VT026005-ZP::GDBD* > *CsChrimson::mVenus* unmated (top) and mated (boaom) females before (left) and during (right) illumination with 10.5 µW/mm^2^ of red light to activate DDAG_B–D neurons. In both cases, photoactivation results in ovipositor extrusion. (F) Average fraction of time unmated and mated females extrude their ovipositor during (red) and before or after (black) three sequential 15-second bouts of photoactivation (*i.e.*, oe index) of the DDAG subtypes labeled by *dsf*^p65AD::Zp^ ∩ *VT026005-ZP::GDBD*. Lights are off for 45 seconds before each bout. Indices are above zero when lights are off because it takes a few seconds for females to fully retract the ovipositor once the photoactivation bout ends. (G) The change in abdominal length upon photoactivation of *dsf*^p65AD::Zp^ ∩ *VT026005-ZP::GDBD* > *CsChrimson::mVenus* in unmated (black) and mated (magenta) females is similar across increasing light intensities. (H) Histogram showing the number of unmated mosaic females in which *CsChrimson::mVenus* was either not expressed (first column) or expressed randomly in DDAG_B–D neurons uni- or bilaterally (columns 2–19) via expression of a heat-inducible Flp recombinase. Most mosaic females expressed *CsChrimson::mVenus* not at all or in one or two DDAG neurons (inset histogram). Upon photoactivation, females performed either no behavior (black bars), an abdominal postural change (orange bars), or an ovipositor extrusion (blue bars). All females within each expression category along the x-axis performed similar behaviors upon photoactivation. (I) Artificial bilateral activation of the DDAG_C or DDAG_D neurons induces ovipositor extrusion. No other neurons were activated in these females. (right) Still-frame images of mosaic *dsf*^p65AD::Zp^ ∩ *VT026005-ZP::GDBD* > *CsChrimson::mVenus* unmated females before (left) and during (right) illumination with 10.5 µW/mm^2^ of red light to activate the DDAG_C or DDAG_D neurons bilaterally and specifically. (left) *CsChrimson::mVenus* expression in the posterior nerve cord of the females shown in the still frames. (D, F, G) show individual points, mean, and SD. A one-way ANOVA Turkey-Kramer test for multiple comparisons was used to measure significance (*P*<0.05). Same leaer indicates no significant difference (*P*>0.05).

We asked how the neurons labeled by *dsf*^p65AD::Zp^ ν *VT026005*-*Zp::GDBD* influence female behavior. During courtship, unmated *D. melanogaster* females signal their willingness to mate by opening their vaginal plates and partially exposing the tube-like ovipositor, a behavior called “vaginal plate opening” or VPO^13,14^. Mated females, however, reject courting males by fully extruding their ovipositor, which may block copulation or male courtship drive^14,15^. The length of the abdomen increases when unmated females open their vaginal plates and mated females extrude their ovipositor, but the change in abdominal length is greater during an ovipositor extrusion than during an opening of the vaginal plates (Figure 2D). We previously showed that transient photoactivation of all DDAG neurons using the mVenus-tagged, red-light-gated cation channelrhodopsin, *CsChrimson::mVenus*^16^, caused unmated females to open their vaginal plates and mated females to extrude their ovipositor^9^ (Figure 2D). Upon photoactivation, *dsf*^p65AD::Zp^ ν *VT026005*-*Zp::GDBD > CsChrimson::mVenus* females fully extruded their ovipositor regardless of their mating status (Figure 2D, E; Video S2, S3). Extrusion of the ovipositor was penetrant and occurred largely during the photoactivation period (Figure 2F), and quantitatively similar behaviors were observed across a range of stimulus intensities (Figure 2G).

We next tested whether activity of the DDAG_B–D neurons was required for ovipositor extrusion in mated females. In addition to labeling a subset of female-specific DDAG neurons, the intersection of *dsf*^p65AD::Zp^ and *VT026005*-*Zp::GDBD* labels neurons in the brain (Figure 2A). To target the DDAG_B–D neurons specifically, we intersected *dsf*^p65AD::Zp^ ν *VT026005*-*Zp::GDBD* with *dsx*^LexA::p65^ (Figure 3A) and used the three-way intersection to suppress the activity of the DDAG_B–D neurons by driving the expression of a GFP-tagged version of the inwardly rectifying K+ channel, *Kir2.1*^17^ (Figure 3B). Unmated females expressing *Kir2.1::gfp* in the DDAG_B–D neurons copulated with males at a rate that was similar to control unmated females (Figure 3C) and opened their vaginal plates during courtship at a frequency comparable to control females (Figure 3D). However, mated *dsf*^p65AD::Zp^ ν *VT026005*-*Zp::GDBD* ν *dsx*^LexA::p65^ *> Kir2.1::gfp* females extruded their ovipositor during courtship with a modest reduction in frequency (Figure 3E) and laid fewer eggs (Figure 3F) compared to controls. Thus, the activity of one or more DDAG subtypes labeled by the *dsf*^p65AD::Zp^ ν *VT026005*-*Zp::GDBD* intersection contribute to ovipositor extrusion and egg laying in mated females. The modest effects on ovipositor extrusion and egg laying frequency from silencing these neurons may suggest the involvement of additional neural circuit elements.

**Figure 3.**
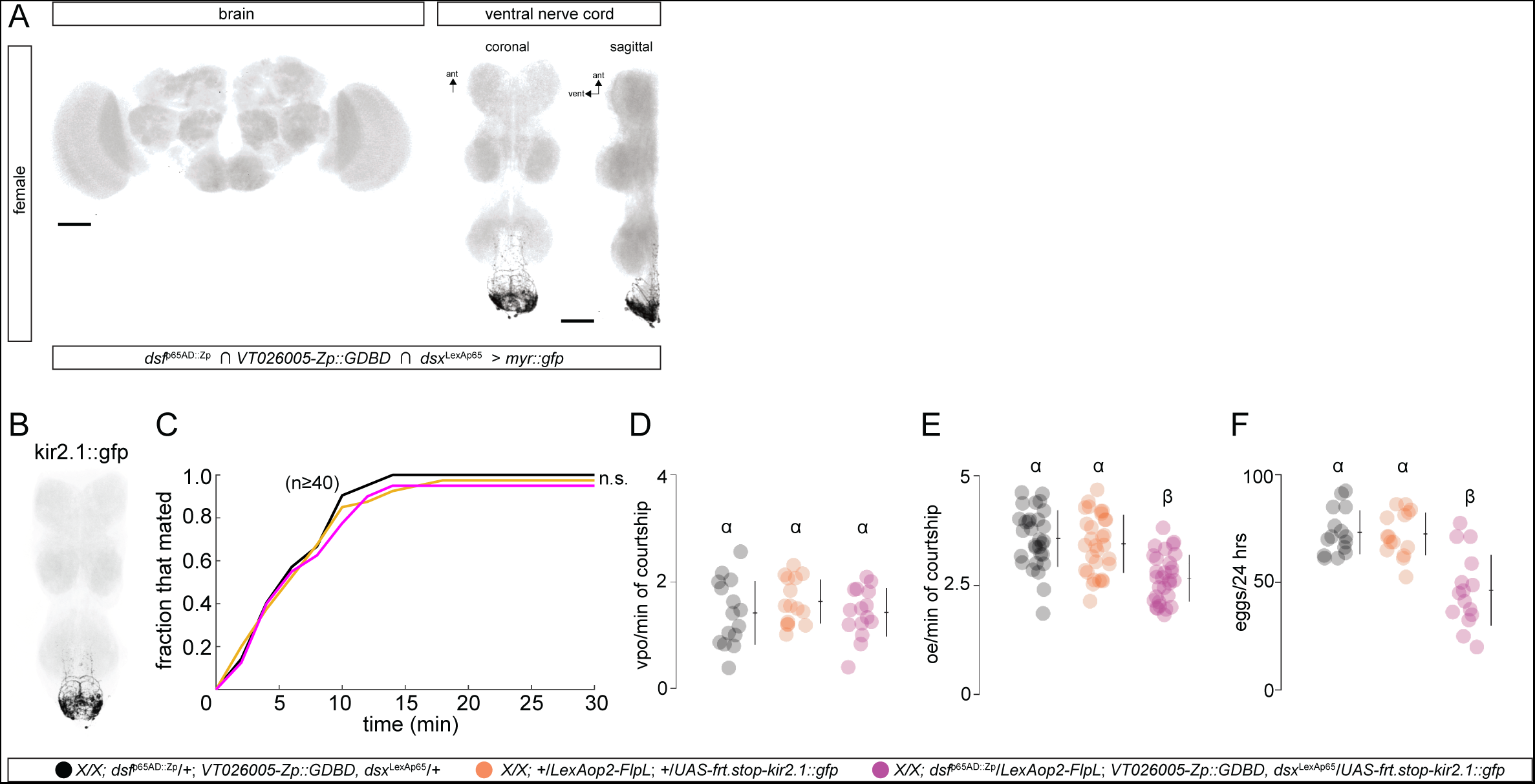
Activity of neurons labeled by *dsf*^p65AD::Zp^ ∩ *VT026005-ZP::GDBD* is necessary for ovipositor extrusion and egg laying in mated females. (A) Confocal images of the brain and VNC from *dsf*^p65AD::Zp^ ∩ *VT026005-ZP::GDBD* ∩ *dsx*^LexA::p65^ > *myr::gfp* females. GFP-expressing neurons are labeled in black and DNCad (neuropil) is shown in light gray. (B) Confocal image of a VNC from *dsf*^p65AD::Zp^ ∩ *VT026005-ZP::GDBD* ∩ *dsx*^LexA::p65^ > *kir2.1::gfp* females. Kir2.1::GFP-expressing neurons are labeled in black and DNCad (neuropil) is shown in light gray. (C, D) Unmated females with inhibited DDAG_B–D neurons using *kir 2.1::gfp* are similarly receptive to male courtship relative to two control genotypes. (C) Fraction of unmated females that mated with a naïve *Canton S* male over a 30-min period. Significance (*P*<0.05) was measured using a Logrank test. n.s. = not significant. (D) Number of times an unmated female opened her vaginal plates per minute of active courtship from a naïve *Canton S* male. (E, F) Ovipositor extrusion and egg laying are reduced in mated females with inhibited DDAG_B–D. (E) Number of times a mated female extruded her ovipositor per minute of active courtship from a naïve *Canton S* male. (F) Number of eggs laid by a mated female 24-hr after mating. (D–F) show individual points, mean, and SD. A one-way ANOVA Turkey-Kramer test for multiple comparisons was used to measure significance (*P*<0.05). Same leaer indicates no significant difference (*P*>0.05).

To determine the specific DDAG neuronal subtype that influences ovipositor extrusion, we randomly expressed *CsChrimson::mVenus* in one or more neurons labeled by the *dsf*^p65AD::Zp^ ν *VT026005*-*Zp::GDBD* intersection using a Flp-based approach. By stochastically expressing a Flp recombinase under the control of a heat-inducible promoter, we randomly excised a transcriptional stop casseae from a UAS-regulated *CsChrimson::mVenus* transgene whose expression was driven by *dsf*^p65AD::Zp^ ν *VT026005*-*Zp::GDBD*. Using this strategy, we generated a population of mosaic females (n=103) that randomly expressed *CsChrimson::mVenus* in one or more of the DDAG neurons labeled by *dsf*^p65AD::Zp^ ν *VT026005*-*Zp::GDBD*, producing 18 distinct groups of mosaic females (Figure 2H). Most of these females expressed *CsChrimson::mVenus* in one or two DDAG neurons (Figure 2H inset). Females were tested in an optogenetic activation experiment before their ventral nerve cords were dissected and stained with antibodies to GFP to reveal the identity of DDAG neurons that expressed the *CsChrimson::mVenus* in each female.

All mosaic females expressing *CsChrimson::mVenus* specifically in a bilateral pair of DDAG_C (n=5) or DDAG_D (n=4) neurons extruded their ovipositor during bouts of photoactivation (Figure 2H, I; Video S4, S5). Unilateral activation of the DDAG_C (n=15) or DDAG_D (n=13) neurons caused a change in abdominal posture with no extrusion of the ovipositor (Figure 2H, Video S6, S7). It is unclear if the postural change is associated with any displacement of the vaginal plates. Photoactivation of a unilateral DDAG_B neuron (n=16) failed to evoke any obvious behavior (Figure 2H, Video S8). We did not obtain mosaic females that expressed *CsChrimson::mVenus* specifically in a bilateral pair of DDAG_B neurons (Figure 2H). However, bilateral activation of the DDAG_B neurons did not modify the postural change induced by photoactivation of a single DDAG_C or DDAG_D neuron (Figure 2H, Video S9). Taken together, these results demonstrate that the DDAG_C and DDAG_D neurons contribute to ovipositor extrusion in mated females. A functional difference between the DDAG_C and DDAG_D neurons was not detected in our experiments. The contribution of the DDAG_B neurons to female behavior is currently unclear.

### The DDAG neurons originate as embryonic-born neurons in the larval ventral nerve cord

In each hemisphere of the late third-instar larval abdominal ganglion, the *dsf*^Gal4^ allele labels two segmentally repeating interneurons from A1 to A8, and four interneurons at the terminal segments (Figure 4A). The number and gross anatomy of these twenty neurons is similar in female and male larvae, in newly hatched larvae, and in larvae aged 24 and 48 hours after hatching (Figure 4A), indicating that the neurons are born during embryogenesis, and that the expression of the *dsf*^Gal4^ allele is stable through larval life. We posited that a subset of these twenty *dsf*-expressing neurons in each hemisphere of the larval abdominal ganglion become the DDAG neurons of adult females and males. We tested this by visualizing *dsf*^Gal4^ and *dsx*^LexA::p65^ expression in the abdominal ganglion of pupae staged at several times during pupal development. From 0 to 18 hours after puparium formation (APF), the twenty *dsf*-expressing neurons we observed in the abdominal ganglion of larvae were identifiable in the ventral nerve cord of female and male pupae (Figure 4B, C). By 18 hours APF, approximately eleven of these neurons had gained *dsx*^LexA::p65^ expression in both sexes (Figure 4B, C). In neuromeres A3–A7, one of the two *dsf*-expressing neurons in each hemisegment was labeled by *dsx*^LexA::p65^, and all six *dsf*-expressing neurons in A8 and the terminal segments co-expressed *dsx*^LexA::p65^ (Figure 4B). The gain of *dsx*^LexA::p65^ expression in *dsf*-expressing neurons occurred gradually and monomorphically in both sexes from 0 to 18 hours APF, but by 36 hours APF, approximately eight *dsf*^Gal4^ and *dsx*^LexA::p65^ co-expressing neurons were absent in males (Figure 4C). We previously demonstrated that the difference in DDAG neuron number between adult females and males was due to *dsx*-dependent apoptosis in males^9^, indicating that the loss of *dsf*^Gal4^ and *dsx*^LexA::p65^ co-expressing neurons in male pupae aged 36 hours APF is due to cell death. At 48 hours APF, approximately eleven and three *dsf*^Gal4^ and *dsx*^LexA::p65^ co-expressing neurons were present in the abdominal ganglion of females and males, respectively, corresponding to the number of DDAG neurons in adults (Figure 4C).

**Figure 4.**
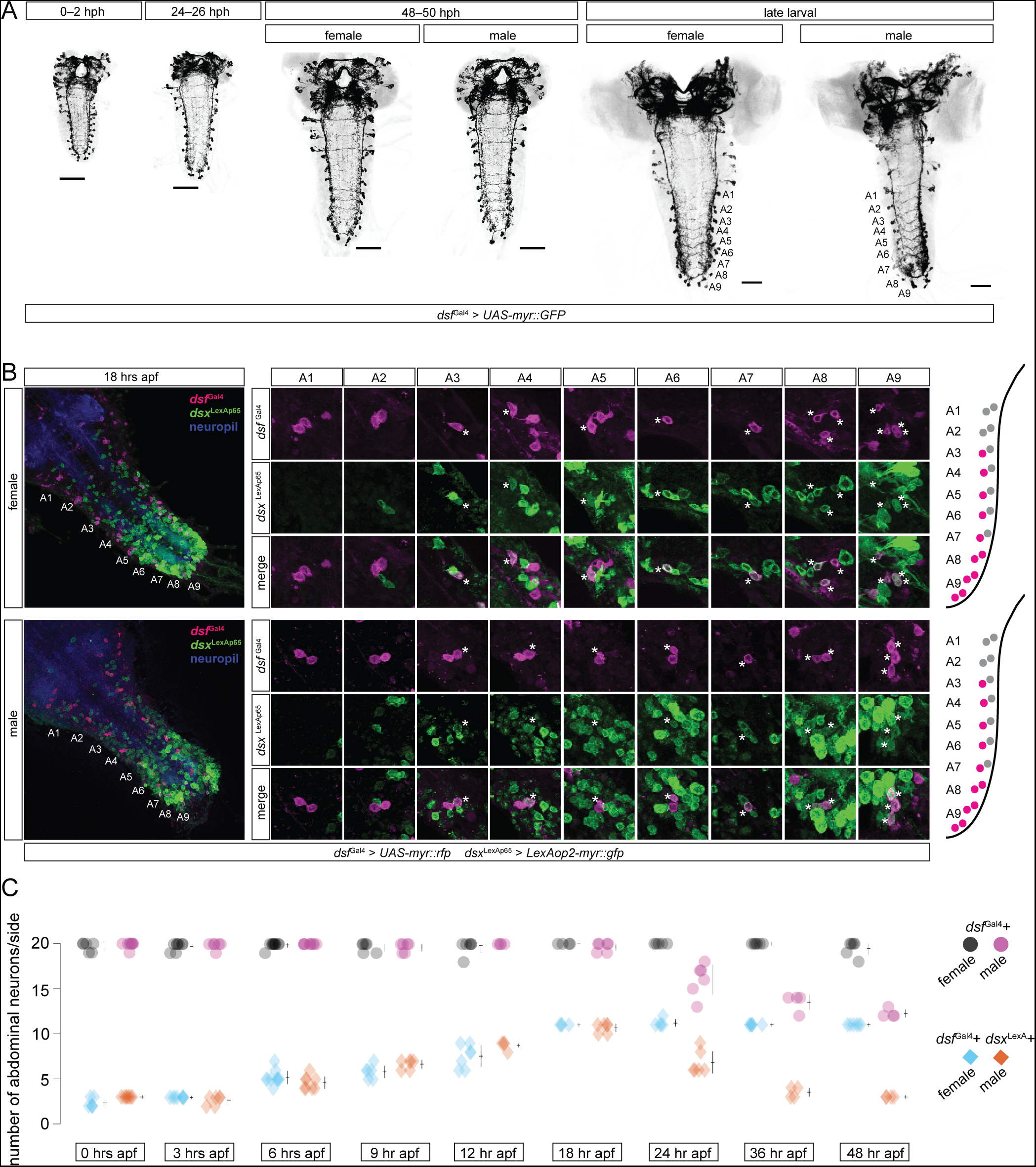
The DDAG neurons originate as embryonic-born *dsf*-expressing interneurons in the abdominal ganglion of larvae. (A) Confocal images of central nervous systems (CNSs) from *dsf*^Gal4^ > *myr::gfp* larvae at various hours post-hatching (hph). GFP-expressing neurons are labeled in black and DNCad (neuropil) is shown in light gray. *dsf*^Gal4^ labels 20 interneurons per hemisphere in the abdominal ganglion of female and male larvae. These neurons are observed in newly hatched larvae and are thus born during embryogenesis. The sex of the larvae at 0–2 and 24–26 hph was not determined. (B) A subset of *dsf*-expressing neurons in the abdominal ganglion of larvae acquire *dsx* expression during pupal life. (left) Confocal images of the VNC from *dsf*^Gal4^ > *myr::rfp*, *dsx*^LexA::p65^ > *myr::gfp* female and male pupae at 18 hours after puparium formation (APF). RFP-expressing neurons are labeled in magenta, GFP-expressing neurons are in green, and DNCad (neuropil) is shown in blue. (right) *dsf*^Gal4^-expressing neurons at each abdominal neuromere are shown. Neurons that co-express *dsx*^LexA::p65^ are labeled with an asterisk. From A3–A7, one of two *dsf*^Gal4^-expressing neurons are also positive for *dsx*^LexA::p65^, whereas all *dsf*^Gal4^-expressing neurons posterior to A7 co-express *dsx*^LexA::p65^. An illustration summarizing the expression of *dsf*^Gal4^ and *dsx*^LexA::p65^ in the abdominal ganglion at 18 hours APF is shown to the right. Cells that co-express *dsf*^Gal4^ and *dsx*^LexA::p65^ are shown in magenta. (C) The number of *dsf*^Gal4^-expressing neurons (circles) and *dsf*^Gal4^-, *dsx*^LexA::p65^-co-expressing (*i.e.*, DDAG) neurons (diamonds) per hermisphere in the abdominal ganglion of females and males is shown. From 0–18 hours APF, both sexes have ~20 *dsf*^Gal4^-expressing neurons, and the number of *dsf*^Gal4^- and *dsx*^LexA::p65^-co-expressing neurons gradually increases to ~11 neurons. Between 18–48 hours APF, ~8 DDAG neurons gradually disappear in males but not in females leaving a total of 11 and 3 DDAG neurons in females and males, respectively. Individual points, mean, and SD are shown.

To further test that the DDAG neurons are derived from *dsf*-expressing neurons in the abdominal ganglion of larvae, we repeated the Flp-based genetic intersection between *dsf*^Gal4^ and *dsx*^LexA::p65^, but this time, we used *UAS-FlpSwitch* to conditionally activate the recombinase in *dsf*^Gal4^-expressing cells only during larval life. Larvae were fed mifepristone, causing the FlpSwitch to excise a transcriptional stop casseae from a LexAop-controlled *myr::gfp* transgene in larval cells labeled by the *dsf*^Gal4^. The *dsx*^LexA::p65^ allele was used to drive the expression of *myr::gfp* in *dsx*-expressing neurons of adults. GFP expression was observed in the DDAG neurons of adults of both sexes (Figure S1), consistent with the notion that the DDAG neurons originate as *dsf*-expressing neurons in the larval abdominal ganglion. We conclude that the entire population of DDAG neurons are embryonic-born *dsf*-expressing neurons that are present in larvae of both sexes. During pupal life, approximately eleven *dsf*-expressing neurons in the abdominal ganglion of both sexes gain *dsx* expression (Figure 4B, C), eight of which are subsequently lost in males due to *dsx*-mediated apoptosis^9^.

### Two segmental homologs of the A26g neuron in larvae become the DDAG_C and DDAG_D neurons of adult females

We next sought to identify the larval counterparts of specific DDAG neuronal subtypes. We found that the intersection of *dsf*^p65AD::Zp^ and *VT026005*-*Zp::GDBD* used above to target the DDAG_B–D subtypes labeled a single, bilateral, and segmentally repeating *dsf*-expressing interneuron in segments A4–A6 of the larval abdominal ganglion of both sexes (Figure 5A, B). Stochastic labeling of individual neurons labeled by the Split-Gal4 demonstrated that these neurons correspond to the A26g interneuron (Figure 5C; J. Truman, personal communication).

**Figure 5.**
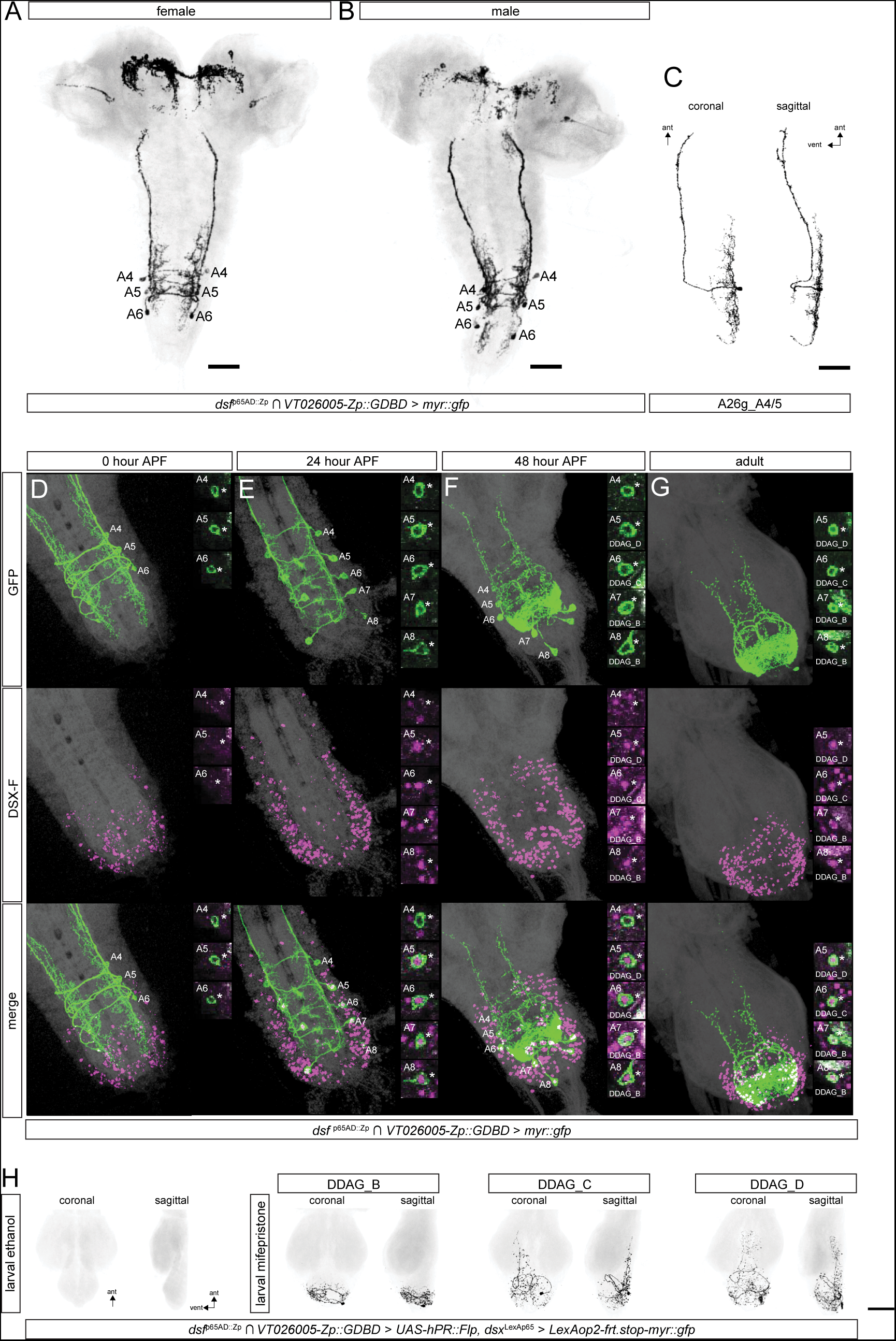
Two A26g neurons in the larval abdominal ganglion become the DDAG_C and DDAG_D neurons of adult females. (A, B) Confocal images of larval CNSs from *dsf*^p65AD::Zp^ ∩ *VT026005-ZP::GDBD* > *myr::gfp* (A) females and (B) males. GFP-expressing neurons are labeled in black and DNCad (neuropil) is shown in light gray. The A26g neuron is labeled in segments A4–A6. (C) A confocal image of a single A26g neuron at A4 or A5 using the multicolor Flp-out technique. Coronal and sagiaal views are shown. (D–G) Confocal images of the abdominal ganglion at various time points after puparium formation (APF) from *dsf*^p65AD::Zp^ ∩ *VT026005-ZP::GDBD* > *myr::gfp* females. GFP-expressing neurons are labeled in green and DSX-F-expressing neurons are labeled in magenta. (D) At 0 hours APF, the A26g neuron at A4–A6 is labeled by GFP, but none of them express DSX-F. (E) At 24 hours APF, the A26g neuron at A5 and A6 express DSX-F. Two additional neurons at A7 and A8 are labeled by *dsf*^p65AD::Zp^ ∩ *VT026005-ZP::GDBD*, both of which express DSX-F. (F) By 48 hours APF, the anatomy of the neurons labeled by *dsf*^p65AD::Zp^ ∩ *VT026005-ZP::GDBD* is similar to that of the DDAG_B–D neurons of adults. (G) By adulthood, only neurons at A5 (DDAG_D), A6 (DDAG_C), A7, and A8 (DDAG_B) are labeled by GFP and DSX-F. (H) Confocal images of posterior VNCs from *dsf*^p65AD::Zp^ ∩ *VT026005-ZP::GDBD* > *UAS-hPR::Flp*, *dsx*^LexAp65^ > *LexAop2-frt.stop-myr::gfp* adult females fed ethanol- or mifepristone-containing food during larval life. GFP-expressing neurons are labeled in black and DNCad (neuropil) is shown in light gray. Expression of GFP in the DDAG_C and DDAG_D neurons of adults indicates that the cells existed among the *dsf*^p65AD::Zp^ ∩ *VT026005-ZP::GDBD*-expressing neurons in the larval abdominal ganglion. Although the Split-Gal4 labels the DDAG_B neurons after larval life (*i.e.*, between 0–24 hours APF), the DDAG_B neurons were also labeled in these experiments. This was likely due to the perdurance of residual mifepristone after pupariation.

The A26g neuron of larvae and the DDAG_C and DDAG_D neurons of adult females display several anatomical similarities (Video S1). All three neurons have a cell body on the dorsal side of the abdominal ganglion, a ventral arch, and a dorsally-located contralateral branch that projects anteriorly. We therefore hypothesized that two segmental homologs of the A26g neuron labeled in larvae by *dsf*^p65AD::Zp^ ν *VT026005*-*Zp::GDBD* metamorphose into the DDAG_C and DDAG_D neurons of adult females. To test this, we first visualized the A26g neurons labeled by *dsf*^p65AD::Zp^ ν *VT026005*-*Zp::GDBD* over the course of pupal development in females and probed DSX-F protein expression. At the onset of pupariation, the A26g neurons at A4–A6 were labeled by *dsf*^p65AD::Zp^ ν *VT026005*-*Zp::GDBD* but none of them co-expressed DSX-F (Figure 5D). By 24 hours APF, however, the A26g neuron at A5 and A6, but not A4, had gained DSX-F expression (Figure 5E). Two additional neurons, one at A7 and one at A8, were labeled by the Split-Gal4 and both were also marked by DSX-F (Figure 5E). By 48 hours APF, the gross morphology of the neurons labeled by *dsf*^p65AD::Zp^ ν *VT026005*-*Zp::GDBD* had transformed to the likeness of the DDAG_B–D neurons of adult females (Figure 5F, G). The cell bodies of the DSX-F-expressing A5 and A6 neurons were positioned on the dorsal side of the abdominal ganglion where the soma of the DDAG_D and DDAG_C neurons are normally located, and the A7 and A8 cell bodies were at the tip of the nerve cord, where the DDAG_B neurons are found. By adulthood, the DSX-F-non-expressing A26g neuron at A4 had disappeared (Figure 5G). The DDAG_D neuron is anterior to the DDAG_C neuron, suggesting that the A26g neuron at segments A5 and A6 transform into the DDAG_D and DDAG_C neurons, respectively. The DDAG_B neurons are likely derived from *dsf*-expressing neurons at A7 and A8 that become marked by *dsf*^p65AD::Zp^ ν *VT026005*-*Zp::GDBD* during pupal development prior to 24 hours APF. The larval counterparts of the DDAG_B neurons are currently not known.

To confirm that the A26g neurons in larvae become the DDAG_C and DDAG_D neurons, we repeated the FlpSwitch-based genetic intersection we described above but between *dsx*^LexA::p65^ and *dsf*^p65AD::Zp^ ν *VT026005*-*Zp::GDBD.* The Split-Gal4 was used to drive the expression of *UAS-FlpSwitch* in the A26g neurons and larvae were fed mifepristone to activate the FlpSwitch during larval life. The FlpSwitch then excised a transcriptional stop casseae from a LexAop-controlled *myr::gfp* transgene driven by *dsx*^LexA::p65^. The DDAG_C and DDAG_D neurons of adult females were labeled by GFP (Figure 5H), further confirming that the DDAG_C and DDAG_D neurons are indeed derived from two segmental homologs of the A26g neurons in larvae.

### DSF and DSX-M regulate the survival of the DDAG_C and DDAG_D neurons

We previously showed that DSF activity is required for the survival of a subset of DDAG neurons in females, whereas DSX-M promotes the cell death of female-specific DDAG neurons in males^9^. We sought to determine how DSF and DSX contribute to the development of the DDAG_C and DDAG_D neurons specifically. The *dsf*^p65AD::Zp^ ν *VT026005*-*Zp::GDBD* intersection was used to visualize the DDAG_C and DDAG_D neurons while driving the expression of a validated UAS-regulated short hairpin/miRNA (ShmiR) targeting *dsf* or *dsx* transcripts. *Dsx* transcripts are sex-specifically spliced and translated to produce female- and male-specific isoforms of DSX proteins^18^. Knock-down of *dsx* transcripts in females did not cause any obvious change in DDAG_C or DDAG_D anatomy (Figure S2A, B), whereas depletion of male-specific *dsx* transcripts caused a gain of DDAG_C and DDAG_D neurons with an arborization paaern like that of wild-type females (Figure S2A’, B’). Female-like DDAG neurons are resurrected in males with the ectopic expression of the cell death inhibitor, P35^9^, suggesting that the gain of the DDAG_C and DDAG_D neurons in *dsf*^p65AD::Zp^ ν *VT026005*-*Zp::GDBD* > *UAS-dsx_ShmiR* males is due to loss of cell death. Optogenetic activation of the resurrected DDAG_C and DDAG_D neurons in *dsf*^p65AD::Zp^ ν *VT026005*-*Zp::GDBD* > *UAS-dsx_ShmiR* males induced an extrusion of the male’s terminalia (Video S10), a behavior reminiscent of ovipositor extrusion in mated females. Consistent with the absence of an anatomical phenotype in their DDAG_C and DDAG_D neurons, photoactivation of *dsf*^p65AD::Zp^ ν *VT026005*-*Zp::GDBD* > *UAS-dsx_ShmiR* females caused an extrusion of the ovipositor (Figure S2G, Video S11).

In contrast, the DDAG_C and DDAG_D neurons were lost in *dsf*^p65AD::Zp^ ν *VT026005*-*Zp::GDBD > UAS-dsf_ShmiR* females (Figure S2A, C) and in females carrying loss-of-function mutations in *dsf* (Figure S2D, E). When *dsf*^p65AD::Zp^ ν *VT026005*-*Zp::GDBD* was used to drive the expression of *UAS-dsf_ShmiR* and *UAS-CsChrimson::mVenus*, females failed to extrude their ovipositor during bouts of photoactivation (Figure S2G, Video S12). To confirm that the loss of the DDAG_C and DDAG_D neurons in *dsf* mutant females was due to cell death, we used *dsf*^Gal4^ to drive the expression of *UAS-P35*. Indeed, blockage of cell death by ectopic expression of P35 in *dsf*-expressing neurons of *dsf* mutant females rescued the DDAG_C and DDAG_D neurons (Figure S2D–F). The rescued DDAG_C and DDAG_D neurons appear to lack contralateral ascending projections, suggesting that *dsf* may contribute to neuronal development beyond acting as a pro-survival factor. Knock-down of *dsf* transcripts in *dsf*^p65AD::Zp^ ν *VT026005*-*Zp::GDBD* > *UAS-dsf_ShmiR* males had no effect on the survival of the DDAG_B–D neurons (Figure S2A’, C’). Taken together, these data suggest that DSF promotes the survival of the DDAG_C and DDAG_D neurons in females, whereas DSX-M promotes their cell death in males. Knock-down of *dsf* transcripts in *dsf*^p65AD::Zp^ ν *VT026005*-*Zp::GDBD > UAS-dsf_ShmiR* larvae had no obvious effect upon the development of the A26g neurons (Figure S2H), suggesting that DSF regulates the survival of the DDAG_C and DDAG_D neurons during pupal development.

## DISCUSSION

Many neurons of the *Drosophila* larval nervous system persist through metamorphosis and contribute to adult neural circuits, yet their contribution to sexually dimorphic adult behaviors is unclear. In this paper, we address this gap with two key findings: First, we identify two interneurons in the abdominal ganglion of adult females, the DDAG_C and DDAG_D neurons, that contribute to ovipositor extrusion, a behavior performed primarily by mated and mature females to reject courting males. The neural circuitry that mediates ovipositor extrusion in mated females has been partially delineated recently^15^. In response to the male’s song, *dsx*-expressing pC2l neurons activate a descending neuron called DNp13 that triggers the motor circuits in the abdominal ganglion for ovipositor extrusion^15^. Interestingly, the ability of DNp13 to induce ovipositor extrusion in mated but not unmated females depends upon mechanosensory input during ovulation^15^. The DDAG_C/D neurons may function downstream of DNp13 and ovulation-sensing mechanosensory neurons, perhaps integrating their inputs, but upstream of motor circuits for ovipositor extrusion.

Photoactivation of all DDAG neurons induces vaginal plate opening in unmated females and ovipositor extrusion in mated females^9^. Activating the DDAG_B–D neurons, however, induces ovipositor extrusion regardless of mating status (Figure 2D). This suggests the possibility that the DDAG_A or DDAG_E subtypes or both integrate mating status to inhibit DDAG_C/D-driven ovipositor extrusion in unmated females. Testing this will require the development of new genetic tools that provide access to the DDAG_A and DDAG_E subtypes.

Second, we find that the DDAG_C/D neurons are present in the larval abdominal ganglion as mature sexually monomorphic neurons corresponding to two segmental homologs of the A26g neuron. The function of A26g in larvae is currently unknown. During metamorphosis, the A26g neurons at segments A5 and A6 acquire *dsx* expression in both sexes and are then remodeled in females to become the DDAG_D and DDAG_C neurons, respectively. In males, expression of DSX-M promotes programmed cell death of the A26g neurons during pupal life^9^, whereas DSF activity is necessary for the survival the A26g neurons in females. The mechanism by which DSF and DSX-M regulate A26g apoptosis is unknown. One possibility is that DSX-M antagonizes DSF function in males, thereby allowing the neurons to die during pupal life.

Our results demonstrate that the neural circuits for courtship behavior in *Drosophila* are constructed in part from sex-non-specific larval neurons that are sexually reprogrammed during metamorphosis for functions in adults (Figure 6). Similar observations have been made in *C. elegans* where sex-specific paaerns of synaptic connectivity in adults develop from neurons in juvenile worms with sexually monomorphic connections^19^. How much of the circuitry for dimorphic courtship behaviors in flies is built from reprogrammed larval neurons? Lineage analyses of neurons with sexual identity in the adult brain and thoracic ganglia have shown that most, if not all, are born during post-embryonic neurogenesis and function only in adults^7,8^. However, this may differ in the abdominal nervous system. The abdominal ganglion of adult flies is largely specialized for functions in reproduction, and indeed the majority of *dsx*-expressing neurons are located in the abdominal ganglion^20–22^. Some of these neurons are specific to adults and are born during larval and pupal life from two terminal abdominal neural stem cells, *i.e.*, neuroblasts, with sex-specific paaerns of neurogenesis^23–25^. But many others, like the DDAG neurons, are likely to be derived from embryonic lineages. In contrast to the brain and thoracic nervous system, the vast majority of neuroblasts in the abdominal neuromeres (*e.g.*, A2–A8) finish producing neurons by the end of embryonic life and add very few adult-specific neurons^23^. The contribution of remodeled larval neurons to the circuits for sexually dimorphic behaviors may thus be relatively greater in the abdominal ganglion than in other regions of the fly nervous system.

**Figure 6.**
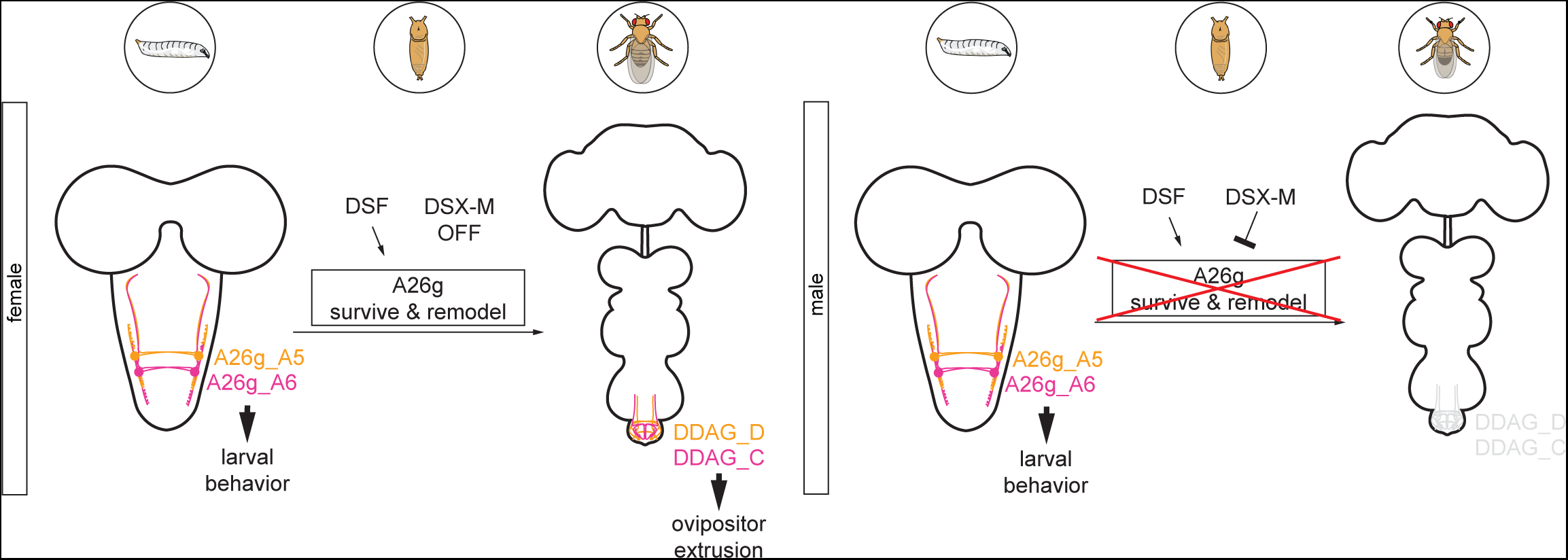
A26g neurons in larvae become the DDAG_D and DDAG_C neurons in females but undergo *dsx*-dependent cell death in males. A model summarizing the results described in this paper. See text for details.

Insect neurons exhibit impressive plasticity as the central nervous system metamorphoses from its larval to adult form. Some larval neurons undergo programmed cell death, but most persist through pupal life to contribute to adult circuits^5^. Larval neurons that persist remodel their axonal and dendritic arbors to regulate similar processes in adults^3,6,26–30^ or trans-differentiate to obtain altogether different functions^3,31^. Courtship and neuronal sexual identity are specific to adults, suggesting that the A26g-to-DDAG_C/D transformation may be a case of trans-differentiation. How the A26g neurons acquire sexual identity and become repurposed during metamorphosis may provide a system to study the regulatory mechanisms underlying developmental reprogramming of the nervous system.

## ACKNOWLEDGEMENTS

We thank J. Truman (University of Washington) for identifying the A26g neuron and suggesting driver lines to target them in larvae; J. Cande and D. Stern (HHMI) for contributing *dsf* alleles; Y. Ding (University of Pennsylvania), J. Lillvis (HHMI), and E. Preger-Ben Noon (Technion) for helpful discussions and comments on the manuscript; and A. McStravog for administrative assistance. T.R.S. is supported by the National Science Foundation (IOS-1845673).

## AUTHOR CONTRIBUTIONS

Conceptualization, T.R.S.; Investigation, J.A.D, J.C.D, K.E.M, M.W., S.L., M.R.M., T.R.S.; Writing – Original Draft, T.R.S., J.A.D.

## STAR METHODS

### KEY RESOURCES TABLE

| REAGENT or RESOURCE | SOURCE | IDENTIFIER |
| --- | --- | --- |
| Antibodies |  |  |
| rabbit anti-GFP | Invitrogen | Cat#A11122; RRID:AB_221569 |
| rat anti-DN-cadherin | Developmental Studies Hybridoma Bank | Cat#DN-Ex#8; RIDD:AB_528121 |
| rabbit anti-DSX-F | Peng et al.<sup>32</sup> | N/A |
| mouse anti-DSX-M | Peng et al.<sup>32</sup> | N/A |
| mouse anti-HA.11 | BioLegend | Cat#MMS-101P; RRID:AB_291261 |
| rat anti-FLAG | Novus Biologicals | Cat#NBP1-06712; RRID:1625981 |
| Fluorescein (FITC)-conjugated donkey anti-rabbit | Jackson ImmunoResearch | Cat#711-095-152; RIDD:AB_2315776 |
| goat anti-rat AlexaFluor 647 | Invitrogen | Cat#A21247; RRID:AB_141778 |
| donkey anti-rat AlexaFluor 568 | Invitrogen | Cat#A78946; RRID:AB_2910653 |
| donkey anti-mouse AlexaFluor 647 | Invitrogen | Cat#A31571; RRID:AB_162542 |
| goat anti-mouse AlexaFluor 488 | Invitrogen | Cat#A32723; RRID:AB_2633275 |
| Chemicals, peptides, and recombinant proteins |  |  |
| DPX | Sigma-Aldrich | Cat#06522 |
| all-*trans*-Retinal | Sigma-Aldrich | Cat#R2500 |
| Mifepristone (RU-486) | Sigma-Aldrich | Cat#475838-50MG |
| Experimental models: Organisms/strains |  |  |
| Canton S | Duckhorn et al.<sup>9</sup> | N/A |
| *w<sup>1118</sup>* | Duckhorn et al.<sup>9</sup> | N/A |
| *dsx<sup>LexA::p65</sup>/TM6B* | Zhou et al.<sup>33</sup> | N/A |
| *dsf<sup>Gal4</sup>/CyO* | Duckhorn et al.<sup>9</sup> | N/A |
| *pJFRC29-10XUAS-IVS-myr::GFP-p10 (attP2)* | Janelia Research Campus (JRC), HHMI | N/A |
| *pJFRC12-10XUAS-IVS-myr::GFP (attP2)* | JRC |  |
| *pJFRC79-8XLexAop-2-FlpL (attP40)* | JRC | N/A |
| *pJFRC41-10XUAS-FRT-STOP-FRT-myr::gfp (su(Hw)attP1)* | JRC | N/A |
| *pJFRC108-20XUAS-IVS-hPR-Flp-p10 (attP2)* | JRC | N/A |
| *pJFRC40-13XLexAop-FRT-STOP-FRT-myr::gfp (attP40)* | JRC | N/A |
| *pJFRC56-10XUAS-FRT-STOP-FRT-kir2.1::gfp (attP2)* | JRC | N/A |
| *pBPhsFlp2::PEST (attP3)* | JRC | N/A |
| *pJFRC201-10XUAS-FRT-STOP-FRT-myr::smGFP-HA (VK0005)* | JRC | N/A |
| *pJFRC240-10XUAS-FRT>STOP>FRTmyr::smGFP-V5-THS-10XUAS-FRT>STOP>FRT-myr::smGFP-FLAG (su(Hw)attP1)* | JRC | N/A |
| *20XUAS-FRT>STOP>FRT-CsChrimson::mVenus (VK5)* | JRC | N/A |
| VT026005-Zp::GDBD (*attP2*) | JRC | N/A |
| UAS-P35<sup>BH1</sup> | BDSC | RRID:BDSC_5072 |
| UAS-dsx_*ShmiR* (*attP2*) | Duckhorn et al.<sup>9</sup> | RRID:BDSC_35645 |
| UAS-dsf_*ShmiR* (*attP2*) | Duckhorn et al.<sup>9</sup> | N/A |
| *dsf*<sup>Del</sup> | Duckhorn et al.<sup>9</sup> | N/A |
| *dsf*<sup>p65AD::Zp/CyO</sup> | This study | N/A |
| *dsf*<sup>DBD::Zp/CyO</sup> | This study | N/A |
| *dsf*<sup>LexA::p65/CyO</sup> | This study | N/A |
| Software and algorithms |  |  |
| Fiji | NIH, USA | https://imagej.net/fiji |
| MATLAB | Mathworks | https://www.mathworks.com/products/matlab.html |

### RESOURCE AVAILABILITY

#### Lead contact

Further information and requests for resources and reagents should be directed to and will be fulfilled by the lead contact, Troy Shirangi.

#### Materials availability

Fly lines generated in this study are available from the lead contact.

#### Data and code availability

All data reported in this paper will be shared by the lead contact upon request. This paper does not report original code. Any additional information required to reanalyze the data reported in this paper is available from the lead contact upon request.

### EXPERIMENTAL MODEL AND SUBJECT DETAILS

*Drosophila melanogaster* stocks were maintained on standard cornmeal and molasses food at 25°C and ~50% humidity in a 12-hr light/dark cycle unless otherwise noted. Fly stocks used in this study are listed in the Key Resources Table. *dsf*^p65AD::Zp^, *dsf*^Zp::GDBD^, and *dsf*^LexA::p65^ alleles were generated using the same strategy as that used to build the *dsf*^Gal4^ allele^9^ except the Gal4 sequence in the donor construct was replaced with sequences encoding *p65AD::Zp*, *Zp::GDBD*, or *LexA::p65*.

### METHOD DETAILS

#### Immunohistochemistry

Nervous systems were dissected in 1X phosphate-buffered saline (PBS), fixed in 4% paraformaldehyde in PBS for 35 minutes, then rinsed and washed in PBT (PBS with 1% Triton X-100). If a blocking step was performed, nervous systems were incubated in 5% normal goat serum or 5% normal donkey serum in PBT for 30 minutes. Tissues were then incubated with primary antibodies diluted in PBT or PBT with block overnight at 4°C. The next day, three washes were performed over the course of several hours before nervous systems were incubated in secondary antibodies diluted in PBT or PBT with block overnight at 4°C. Tissues were then washed three times over the course of several hours and placed on cover slips coated in poly-lysine, dehydrated in an increasing ethanol concentration series, and cleared in a xylene series. Nervous systems were mounted onto slides using DPX mounting medium and imaged on a Leica TCS SP8 Confocal Microscope at 40X magnification. For MultiColor FlpOut experiments, vials containing larvae aged between first and second instars (or 3–4-day old adults) were placed in a 37°C water bath for 1–10 minutes then dissected 2 days later. To stochastically label the DDAG neurons, male and female adults (deprived of food overnight) were placed in vials with food containing 100mM mifepristone (RU-486; Sigma 475838-50MG) for 2 hours and kept in darkness. Flies were then transferred back to vials containing untreated food for 3 days before their VNCs were dissected for staining. For immortalization experiments using mifepristone, three days after crosses were set-up on normal food, 60 µL of 100mM RU-486 was added directly to the food and larvae were raised in darkness on RU-486-treated food until pupariation. Pupae were collected and transferred to vials containing untreated food before eclosure. Control groups were also kept in darkness but were treated with 60 µL of ethanol instead of RU-486. Adult nervous systems were dissected in PBS. The following primary antibodies were used: rabbit anti-GFP (Invitrogen #A11122; 1:1000), rat anti-DN-cadherin (DN-Ex#8, Developmental Studies Hybridoma Bank; 1:50), anti-DSX-F (1:200)^32^, mouse anti-HA.11 (BioLegend #MMS-101P; 1:250), and rat anti-FLAG (Novus Biologicals #NBP1-06712; 1:200). The following secondary antibodies were used: Fluorescein (FITC) conjugated donkey anti-rabbit (Jackson ImmunoResearch #711-095-152; 1:500), AF-647 goat anti-rat (Invitrogen #A21247; 1:500), AF-568 donkey anti-rat (Invitrogen #A78946; 1:500), AF-647 donkey anti-mouse (Invitrogen #A31571; 1:500), AF-488 goat anti-mouse (Invitrogen #A32723), AF-647 goat anti-rat (Invitrogen #A21247; 1:500).

#### Optogenetic assays

Unmated females used in optogenetic assays were raised in darkness and on food containing 0.2mM all-trans-retinal (sigma-Aldrich #R2500) and were incubated at 25°C and ~50% humidity. Once collected, unmated females were grouped in vials consisting of 15–20 flies for 8–12 days before testing. Flies were anesthetized on ice for ~2 minutes, decapitated under low-intensity light, and were given 15–20 minutes to recover before being transferred to individual behavioral chambers (diameter: 10 mm, height: 3 mm). A FLIR Blackfly S USB3, BFS-U3-31S4M-C camera with a 800 nm long-pass filter (Thorlabs, FEL0800) was used to record optogenetic videos in SpinView. Upon testing, chambers were placed on top of an LED panel with continuous infrared (850 nm) light and recurring photoactivating red (635 nm) light using an Arduino script. To measure change in abdominal length before and during vaginal plate opening or ovipositor extrusion, a ruler (cm/mm) was included in the frame to set the scale. The change in abdominal length was calculated as the difference in the abdominal length from the base of the scutellum to the tip of the abdomen before and during photoactivation. Behavior indices were measured by calculating the average fraction of time spent preforming the behavior during the first three 15-second lights-on periods and the first three 45-second light-off periods. For stochastic optogenetic activation, unmated females carrying *hs-Flp*, *UAS-frt.stop-CsChrimson::mVenus*, and the Split-Gal4 were reared on retinal-containing food, grouped in vials consisting of 15–20 flies, and aged for ~3 days before being placed in a 37°C water bath for periods ranging from 20–60 minutes. Females were then transferred to new vials containing retinal food and aged for an additional 5 days before being tested in an optogenetic activation experiment as described above. Following optogenetics, the VNC of each female was dissected and placed singly in wells of a 60-well mini tray (Fischer Scientific #12-565-155) for staining. Each VNC was subsequently mapped to the female in the optogenetics experiment from which the VNC was obtained.

#### Behavioral assays

Unmated females and males were collected were under CO_2_ and aged for 7–10 days in a 12-hour light/dark cycle and incubated at 25°C and ~50% humidity. Unmated females were group-housed in vials consisting of 15–20 flies, and *Canton S* males were individually housed. Courtship assays were done within the first two hours of the subjective day. Unmated females and *Canton S* males were transferred to individual behavioral chambers (diameter: 10 mm, height: 3 mm) and recorded for 30 minutes using a Sony Vixia HFR700 video camera at 25°C under white light. For experiments using mated females, unmated females were housed with males for 24 hours, anesthetized on ice for ~2 minutes, and mated females were collected into a new vial and given 30 minutes to recover. Mated females and *Canton S* males were loaded to chambers and recorded as described above. Courtship index was measured as the total time the male preformed courtship behaviors divided by the total recording time. Courtship index was measured as the total time the male performed courtship behaviors divided by the observation time which was usually about 5 minutes. Vaginal plate opening (vpo) and ovipositor extrusion (oe) frequency was measured as the total number of times a female performed a vpo or oe in a 6-min period of active male courtship. Egg laying was measured by allowing females to mate with males before transferring them to individual vials to for 24 hours. The total number of eggs laid in 24 hours by each female was then counted.

### QUANTIFICATION AND STATISTICAL ANALYSIS

Data were analyzed using one-way ANOVA with Tukey-Kramer tests for multiple comparisons, Rank Sum tests, or Logrank tests. All p-values were measured in MATLAB.

## Legends for Supplemental Videos

**Video S1. Anatomy of the DDAG_B, DDAG_C, DDAG_D, and A26g neurons, Related to** **Figure 2**.

**Video S2. Optogenetic activation of the DDAG_B–D neurons in unmated females, Related to** **Figure 2**.

**Video S3. Optogenetic activation of the DDAG_B–D neurons in mated females, Related to** **Figure 2**.

**Video S4. Bilateral optogenetic activation of the DDAG_C neurons in unmated females, Related to** **Figure 2**.

**Video S5. Bilateral optogenetic activation of the DDAG_D neurons in unmated females, Related to** **Figure 2**.

**Video S6. Unilateral optogenetic activation of a DDAG_C neuron in unmated females, Related to** **Figure 2**.

**Video S7. Unilateral optogenetic activation of a DDAG_D neuron in unmated females, Related to** **Figure 2**.

**Video S8. Unilateral optogenetic activation of a DDAG_B neuron in unmated females, Related to** **Figure 2**.

**Video S9. Bilateral optogenetic activation of the DDAG_B neurons in unmated females, Related to** **Figure 2**.

**Video S10. Optogenetic activation of a *dsf*^p65AD::Zp^ ∩ *VT026005-Zp::GDBD* > *CsChrimson::mVenus* male with depleted *dsx* transcripts, Related to** **Figure 5**.

**Video S11. Optogenetic activation of a *dsf*^p65AD::Zp^ ∩ *VT026005-Zp::GDBD* > *CsChrimson::mVenus* unmated female with depleted *dsx* transcripts, Related to** **Figure 5**.

**Video S12. Optogenetic activation of a *dsf*^p65AD::Zp^ ∩ *VT026005-Zp::GDBD* > *CsChrimson::mVenus* unmated female with depleted *dsf* transcripts, Related to** **Figure 5**.

