## Supplemental Data 1 for "Developmental remodeling repurposes larval neurons for sexual behaviors in adult *Drosophila*"

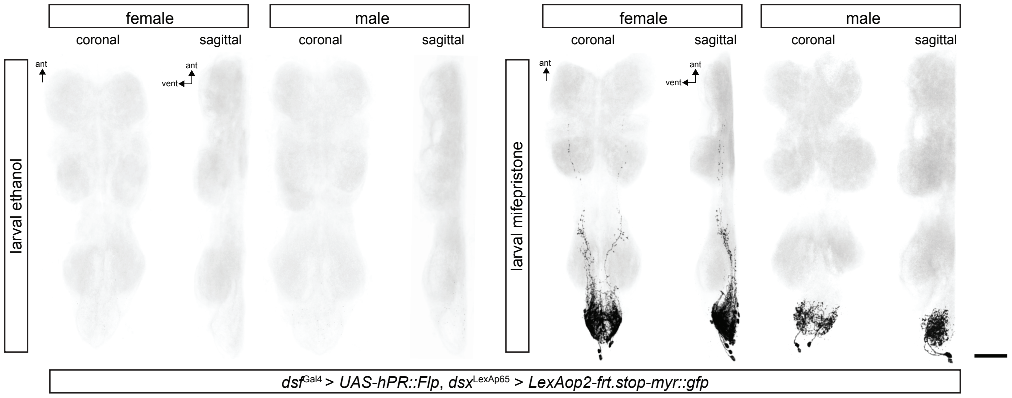


**Figure S1, Related to Figure 4.** Confocal images of VNCs from *dsf*$^{Gal4}$/*LexAop2-frt.stop-myr::gfp*; *dsx*$^{LexA::p65}$/*UAS-hPR::Flp* females and males fed ethanol- or mifepristone-spiked food only during larval life. GFP-expressing neurons are labeled in black and DNCad (neuropil) is shown in light gray. Scale bar = 50 µm.

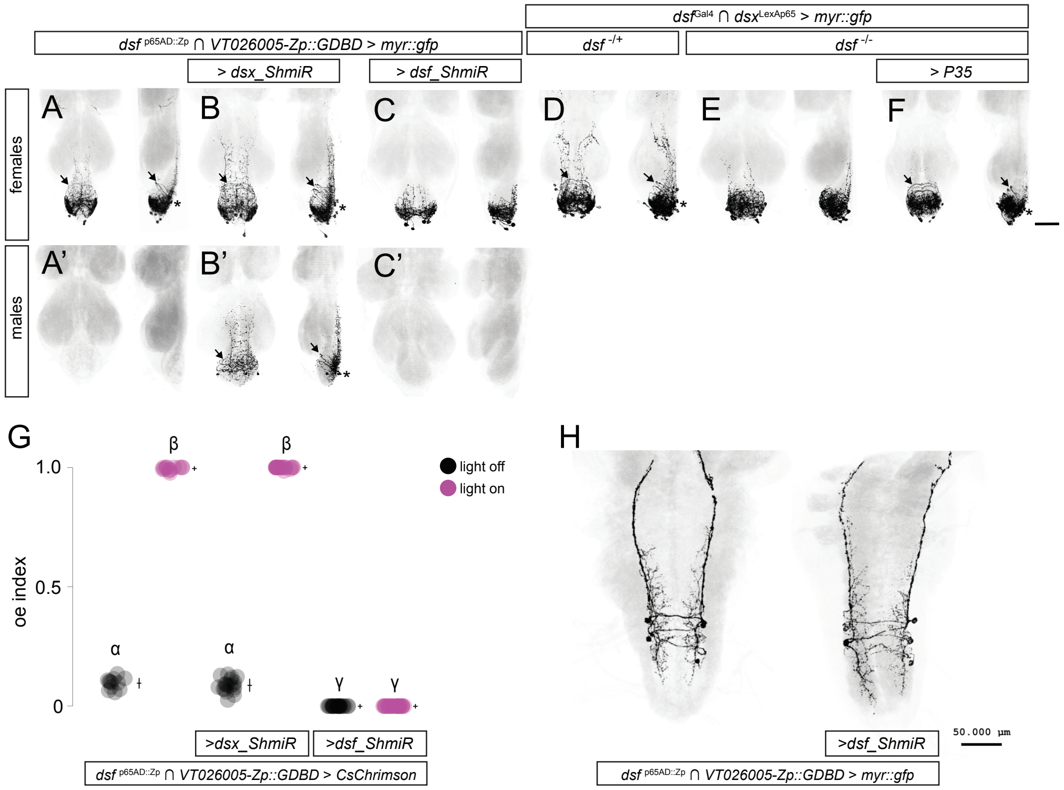


**Figure S2, Related to Figure 5.** Confocal images of posterior VNCs from *dsf*$^{p65AD::Zp}$ ∩ *VT026005-Zp::GDBD > myr::gfp* females (A–D) and males (A'–C'). Control flies (A, A'), and flies carrying *UAS-dsx_ShmiR* (B, B') or *UAS-dsf_ShmiR* (C, C') transgenes are shown. (D–F) Confocal images of posterior VNCs from females. (D) Control flies (*dsf*$^{Gal4}$/*LexAop2-FlpL*; *dsx*$^{LexA::p65}$/*UAS-frt.stop-myr::gfp*), (E) *dsf* mutant flies (*dsf*$^{Gal4}$/*dsf*$^{Del}$, *LexAop2-FlpL*; *dsx*$^{LexA::p65}$/*UAS-frt.stop-myr::gfp*), and (F) *dsf* mutant + p35 flies (*dsf*$^{Gal4}$, *UAS-p35*/*dsf*$^{Del}$, *LexAop2-FlpL*; *dsx*$^{LexA::p65}$/*UAS-frt.stop-myr::gfp, UAS-dsf_ShmiR*) are shown. Coronal and sagittal views of each VNC is shown. Arrows point to the ventral arch of DDAG_C/D neurons. Asterisks note the location of dorsal cell bodies corresponding to the DDAG_C/D neurons. We note that the anterior projections of the DDAG_C/D neurons is absent in (F), suggesting that DSF may influence other aspects of DDAG_C/D development beyond promoting neuronal survival. GFP-expressing neurons are shown in black and NCad (neuropil) is shown in gray. (G) Fraction of time *dsf*$^{p65AD::Zp}$ ∩ *VT026005-Zp::GDBD* unmated females extrude their ovipositor during a 15-sec bout of photoactivation (*i.e.*, oe index). Control females and females carrying *UAS-dsx_ShmiR* or *UAS-dsf_ShmiR* transgenes are shown. Individual points, mean, and SD are shown. A one-way ANOVA Turkey-Kramer test for multiple comparisons was used to measure significance ($P<0.05$). Same letter indicates no significant difference. (H) Confocal images of the VNC from *dsf*$^{p65AD::Zp}$ ∩ *VT026005-Zp::GDBD > myr::gfp* larvae. Knock-down of *dsf* transcripts has no effect on the survival of the A26g neurons suggesting that dsf acts as prosurvival factor during pupal life. Control larvae and larvae carrying a *UAS-dsf_ShmiR* transgene are shown.
